# Lipolysis of host triacylglyceride-rich lipoproteins creates a toxic microenvironment for *Staphylococcus aureus*

**DOI:** 10.64898/2026.01.12.699058

**Authors:** Trushali Vansadia, Camron N. Collins, Anisha Neupane, Nathaniel J. Torres, Ahsan Hameed, Andrew J. Morris, Karen E. Beenken, Mark S. Smeltzer, Ryan M. Allen

**Affiliations:** Department of Physiology and Cell Biology, University of Arkansas for Medical Sciences, Little Rock, AR 72205; University of Arkansas for Medical Sciences, Department of Microbiology and Immunology, Little Rock, AR 72205; Department of Pediatrics, University of Arkansas for Medical Sciences, Little Rock, Arkansas 72205; Arkansas Children’s Nutrition Research Center, Little Rock, AR 72205; Arkansas Children’s Nutrition Research Center and Central Arkansas Veterans Affairs Healthcare System, Little Rock, AR 72205; Department of Microbiology and Immunology, University of Arkansas for Medical Sciences, Little Rock, AR 72205; Department of Biochemistry and Molecular Biology, University of Arkansas for Medical Sciences, Little Rock, AR 72205; Department of Microbiology and Immunology, University of Arkansas for Medical Sciences, Little Rock, AR 72205; Department of Orthopaedic Surgery, University of Arkansas for Medical Sciences, Little Rock, AR 72205; Department of Physiology and Cell Biology, University of Arkansas for Medical Sciences, Little Rock, AR 72205; Winthrop P. Rockefeller Cancer Institute, Little Rock, AR 72205

**Keywords:** *Staphylococcus aureus*, osteomyelitis, lipase, lipoprotein, *agr*, quorum-sensing, triacylglyceride-rich lipoprotein (TRL), low-density lipoprotein (LDL), community-acquired methicillin resistant *S. aureus* (CA-MRSA), lipoprotein lipase (LPL), glycerol ester hydrolase (*geh*), polyunsaturated fatty acids (PUFA)

## Abstract

*Staphylococcus aureus* secretes lipases that hydrolyze glycerol esters to mitigate host detection and scavenge fatty acids for membrane synthesis, but exposure to host-derived polyunsaturated fatty acids (PUFA) can be toxic. Here, we show that host hyperlipidemia protects against *S. aureus* osteomyelitis by engaging this vulnerability. Using a mouse model of severe hyperlipidemia, we show that PUFA-enriched plasma triacylglycerides are associated with reduced bacterial burden and bone destruction from injury-associated osteomyelitis. *In vitro*, human triacylglyceride-rich lipoproteins (TRL) exhibit potent bactericidal activity against post-exponential *S. aureus* that requires lipase activity of glycerol ester hydrolase (*geh*). Pre-exponential exposure to TRL suppressed *agr* quorum-sensing, blunting *geh* expression and toxicity; however, bactericidal activity of TRL could be restored by exogenous lipoprotein lipase. Together, these findings reveal TRL as active antimicrobial particles whose toxicity is unmasked by lipolysis and suggest that modulation of host lipid metabolism may provide new therapeutic opportunities for chronic *S. aureus* infection.

## INTRODUCTION

*Staphylococcus aureus* is a common Gram-positive pathobiont associated with high morbidity and mortality^1^. Its exceptional ability to adapt to diverse tissue microenvironments enables persistence in the bloodstream and colonization of nearly all tissues, including heart and bone^2^. During infection, *S. aureus* deploys an extensive repertoire of virulence factors that disrupt host defenses, promote intracellular survival, and facilitate survival within necrotic^1,3–5^. These traits underlie the severity of many community-acquired infections and contribute to the substantial burden of trauma- and hospital-associated infections.

*S. aureus* is the most common etiological agent of skeletal infections, where it readily forms biofilms that restrict antimicrobial penetration and immune cell access^3,6,7^. Infections of bone (termed herein as osteomyelitis) frequently fails to resolve with systemic antibiotics, even when the causative strain is antibiotic sensitive, and it is common for patients with healthy immune systems to fail to mount effective responses. Rather, treatment of staphylococcal osteomyelitis often requires surgical debridement of infected tissue and long courses of antibiotics^8–10^. Despite these invasive and aggressive interventions, many patients go on to develop chronic infections, suffer disease-related comorbidities, and/or require amputation. These challenges underscore the urgent need for therapeutic innovations that improve infection control and provide effective public health interventions.

Host metabolism is increasingly recognized as a determinant of infection outcome, with lipid metabolism emerging as a key regulator of host-pathogen interactions^11,12^. *S. aureus* can scavenge host-derived fatty acids to support membrane biosynthesis and conserve energy^13–15^. Pathogen uptake of fatty acids is likely passive, and incorporation of fatty acids to membrane lipid pools is mediated by the fatty acid kinase system, in which FakA partners with FakB isoforms that preferentially incorporate saturated (FakB1) or unsaturated (FakB2) fatty acids^16,17^. Recent studies have uncovered a patent fatty acid dehydrogenase system (*FadXDEBA*) capable of β-oxidation in the absence of glucose^18,19^. To access host lipid pools, *S. aureus* secretes broad-specificity lipases capable of hydrolyzing triacylglycerides, phospholipids, and esterified cholesterol^20–23^.

This metabolic flexibility becomes a liability in lipid-rich environments containing antimicrobial fatty acids. For instance, the epidermis is enriched with polyunsaturated fatty acids (PUFAs) and other bactericidal or bacteriostatic lipids derived from sebum^24–27^. Staphylococcal uptake of PUFAs can result in oxidative stress and incorporation of PUFA to phospholipids compromises membrane structure and signaling^28–32^. Lipase activity is therefore tightly regulated, and isolates from deep-tissue infections exhibit higher lipase expression than superficial isolates^33^, suggesting that selective pressure against lipase production varies across host niches.

In deep tissues, extracellular lipids are primarily transported on circulating lipoproteins–high-density lipoprotein (HDL), low-density lipoprotein (LDL) and triacylglyceride-rich lipoproteins (TRL)–which transport esterified fatty acids as neutral lipids and phospholipids between tissues^34,35^. *S.* aureus can scavenge fatty acids from esterified cholesterol on HDL and LDL to support growth in blood^36^, and the host response to bacteremia often features hepatic secretion of serum amyloid A to destabilize HDL and rapidly lower plasma cholesterol^37–39^. In contrast, plasma triacylglycerides (TAG) often rise during acute infection due to enhanced TRL secretion from the liver and reduced lipoprotein lipase (LPL) activity in the vasculature, which slows fatty acid release and particle clearance^40^. Apolipoprotein-B (ApoB), the essential peptide scaffold of LDL and TRL, has been shown to bind auto-inducing peptide (AIP) and suppress virulence factor production induced by the accessory gene regulator (*agr*) quorum-sensing system^41–43^, contributing to improved host responses. Independently, ApoB has also been shown to participate in pathogen immune evasion through binding and sequestration of lipoteichoic acid^44^. It remains unclear whether elevated ApoB-containing lipoproteins benefit the host during chronic *S. aureus* infection.

Here, we show that severe hyperlipidemia in wild-type mice protects against chronic osteomyelitis caused by an aggressive isolate of community-acquired methicillin-resistant *S. aureus* (CA-MRSA). Using human lipoproteins *in vitro*, we demonstrate that TRL become bactericidal following lipolysis by the secreted *S. aureus* lipase glycerol ester hydrolase (Geh) or by recombinant host LPL. We further show that suppression of *agr* by TRL limits *geh* expression and toxicity, revealing a dynamic host-pathogen interaction in which lipid metabolism shapes bacterial survival. These findings identify TRL as active antimicrobial particles and suggest that modulation of host lipid secretion and catabolism may provide new therapeutic avenues for chronic *S. aureus* infection.

## RESULTS

### Host hyperlipidemia antagonizes S. aureus osteomyelitis

To better understand the relationship between host hyperlipidemia and *S. aureus* osteomyelitis we first transduced wild-type mice with an adeno-associated vector (AAV) to over-express a gain-of-function mutant of proprotein-convertase subtilisin/kexin-9 (PCSK9) that causes LDL-independent degradation of LDLR and effectively abolishes ApoB clearance^45^. Thereafter, mice were chronically fed a Western diet (WD) to induce hyperlipidemia (**Figure 1A**). As controls, wild-type mice were transduced with an AAV overexpressing GFP and fed a chow diet (CD). After 14 weeks of AAV/diet treatment, mice in each cage were randomized to receive a traumatic injury of the femur that was subsequently inoculated with either 1 x 10^6^ CFU of community-acquired methicillin-resistant *S. aureus* (CA-MRSA) USA300 lineage strain L.A. County (LAC) or sterile saline (sham)^46^. Wounds were sutured and chronic osteomyelitis was allowed to proceed for two weeks prior to harvest. As expected, mice transduced with AAV-PCSK9 and fed WD had marked hypercholesterolemia relative to AAV-GFP/CD control mice (**Figure 1B**). In AAV-GFP/CD mice, no change in total plasma cholesterol was observed between sham and infected mice and minimal changes in TRL, LDL or ApoB as determined by size-exclusion chromatography (SEC, fractions 14-15; **Fig. 1C; Supplemental Figure S1**). In contrast, MRSA infection in the AAV-PCSK9/WD group was associated with a reduction of total cholesterol, and cholesterol ApoB, and Apolipoprotein-E associated with TRL-, remnant- and LDL-sized particles were also reduced (**Figure 1B,C; Supplemental Figure S1**). As we have previously reported that host lipoproteins can traffic microbial sRNA, we also measured the distribution of RNA across SEC-fractionated plasma (**Figure S1**)^47–49^. Intriguingly, MRSA infection was associated with increased RNA in fractions corresponding with albumin (fraction 20-21), and to a lesser extent, HDL as defined by Apo-AI (fractions 18-19; **Supplemental Figure S1**). Expansion of the LDL pool by AAV-PCSK9/WD was associated with a modest increase in RNA associated with LDL and TRL (fractions 9-16), but this was not altered by infection. Consistent with reduced plasma cholesterol in the MRSA group, we also observed a modest reduction in atherosclerotic plaque burden in the aortic sinus relative to sham AAV-PCSK9/WD mice (**Supplemental Figure S2**). We theorize that reduced lipids and plaque burden are due to reduced food consumption in infected mice, although this could not be directly measured in our mixed caging strategy. This rationale is supported by the fact that hyperlipidemia and plaque development in this model is dependent on consumption of WD, and dietary intervention is a common method to induce plaque regression in short periods of timef^50,51^.

**Figure 1.**
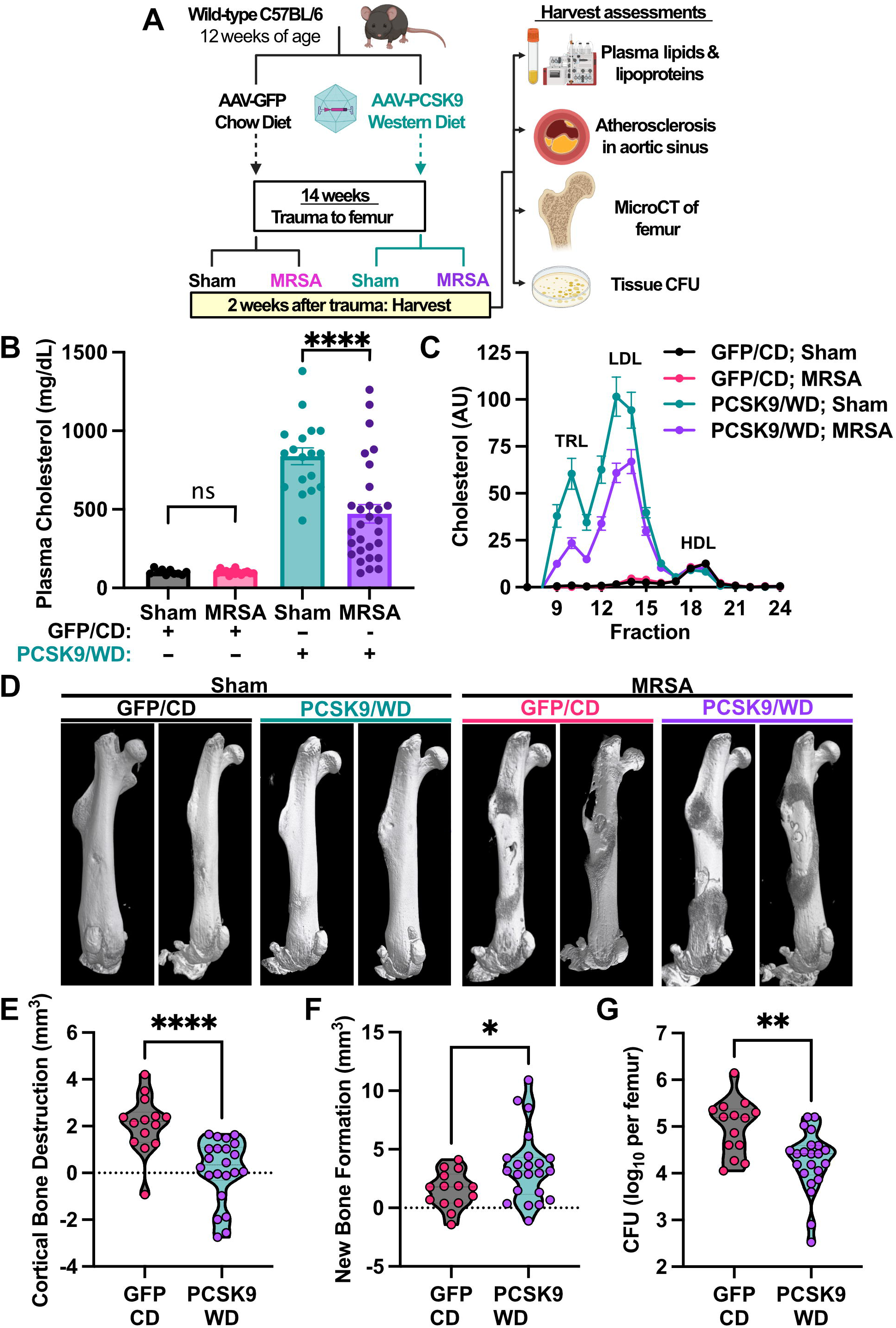
Host hyperlipidemia antagonizes S. aureus osteomyelitis. **A**) Schematic depicting the *in vivo* study design and endpoints. **B**) Plasma cholesterol levels at harvest of mice transduced with AAV-GFP and fed the chow diet (CD) or transduced with AAV-PCSK9 and fed the western diet (WD). Each cage contained mice randomized to sham or MRSA osteomyelitis treatments: AAV-GFP/CD+Sham, n=12; AAV-GFP/CD+MRSA, n=17; AAV-PCSK9/WD+Sham, n=19; AAV-PCSK9/WD+MRSA, n=29. Two-way ANOVA with Sidak’s post-hoc test, ****P_adj_ <0.0001. **C**) Plasma was fractionated by size-exclusion chromatography and total cholesterol in each fraction was determined by colorimetric kit. Major peaks are denoted as triacylglyceride-rich lipoprotein (TRL), low-density lipoprotein (LDL), and high-density lipoprotein (HDL). **D**) Representative μCT reconstructions of femurs from each treatment group. Note: 3 of 17 AAV-GFP/CD+MRSA femurs and 7 of 29 AAV-PCSK9/WD+MRSA femurs were broken during harvest and thus excluded from bacterial burden and μCT analysis. **E)** Bacterial burden of intact femurs infected with MRSA; CFU = Colony forming units. **F)** Quantitative analysis of cortical bone destruction and **G)** new bone formation in intact femurs as determined by μCT analysis. (E-G) Mann-Whitney U-test; *P<0.05, **P<0.01, ****P<0.0001. Data are mean +/- standard error of the mean (SEM).

We next evaluated the impact of hyperlipidemia on bone disease using micro-computed tomography (μCT; **Figure 1D-F**). A significant reduction in cortical bone destruction and a significant increase in new bone formation was observed in hyperlipidemic mice relative to controls. Consistent with less bone destruction, we also observed a significant reduction in bacterial burden in femurs from hyperlipidemic mice (**Figure 1G**). No CFU were observed in blood of either treatment group, nor was there gross evidence of dissemination to ascending lymph nodes of any mice. Infection significantly increased hepatic expression of some acute phase proteins (*Saa1*, *Saa2,* and *Lcn2*), but not others (*A1bg, A2m*, and *Hamp*; **Supplemental Figure S3**). Hepatic expression of apolipoproteins *Apoa1*, *Apob*, and *Apoe* were not significantly altered by diet or *S. aureus* osteomyelitis, nor was *Pcsk9*. Hepatic expression of the pro-inflammatory cytokine *Tnf* was increased by diet, but not infection, whereas *Il6* expression was repressed in MRSA-infected mice transduced with AAV-PCSK9-D377Y relative to infected mice transduced with AAV-GFP. Taken together, these data suggest that increased circulation of host lipoproteins may benefit the host response to *S. aureus* osteomyelitis.

### Western diet feeding increases circulating PUFA in plasma TAG

As LDL has been shown to support *S. aureus* growth in blood by donating fatty acids from esterified cholesterol to the pathogen^36^, we were surprised that high LDL-C in AAV-PCSK9 mice fed WD was associated with reductions in bacterial burden and cortical bone destruction relative to controls (**Figure 1D-F**). As TRL have a central function in distributing fatty acids as inert TAG throughout the body, we hypothesized that increased TRL in AAV-PCSK9 mice fed WD may facilitate enhanced delivery of PUFA to an infected wound and may contribute to reduced bacterial burden and bone disease in these mice. Indeed, AAV-PCSK9/WD treatment increased plasma TAG levels 12-fold in sham mice and 6-fold in mice with MRSA osteomyelitis (**Figure 2A**). Although acute infection often results in sustained increases in plasma TAG in humans, we observed a trend towards reduced TAG in MRSA infected AAV-GFP mice fed the chow diet, and a 2-fold reduction in plasma TAG in MRSA infected AAV-PCSK9 mice relative to sham controls. However, these results are consistent with minimal changes in acute phase response genes in the liver (**Figure S2**). TAG were enriched with large TRL particles as determined by SEC (fractions 9-11), but TAG was also found on remnant- and LDL-sized particles (fractions 12-15; **Figure 2B**). Signal detected with HDL and albumin (fractions 18-21) likely reflects mono- and di-acylglycerides derived from lipase activity (e.g., endothelial lipase, hepatic lipase) on glycerol esters bound to these particles/proteins - products that are indistinguishable from TAG in the enzymatic assay. To evaluate the fatty acid profile of TAG, we analyzed whole plasma using untargeted high-resolution mass spectrometry-based lipidomics and identified 161 unique TAG species (**Supplemental Table S1**). For all treatment groups, TAG primarily contained a mixture of saturated fatty acids and monounsaturated fatty acids, which is consistent with the low PUFA content of milkfat source of WD (<5% PUFA). There was an approximately 3-fold increase in the percentage of TAG containing PUFA in AAV-PCSK9 mice fed WD relative to control mice fed the chow diet (**Figure 2C**). This enrichment in the AAV-PCSK9/WD groups was predominantly associated with TAGs containing fatty acids with >4 double bonds (**Figure 2D**). For example, 52:10 TAG species were enriched more than 50-fold in PCSK9/WD mice (**Figure 2E)**, and 48:4 TAG containing arachidonic acid (20:4) was found nearly exclusively in PCSK9/WD mice (**Figure 2F**). Conversely, TAG with only two or three double bonds were relatively enriched in GFP/SCD mice. Several TAG species were significantly reduced in MRSA-infected mice relative to sham mice in both treatment groups (e.g., 49:4, 48:3, and 48:2). Although the WD used is not an enriched source of dietary PUFA, we speculate that increased circulating TRL mass and relative enrichment of several TAG containing PUFA may contribute to improved host defense against staphylococcal osteomyelitis.

**Figure 2.**
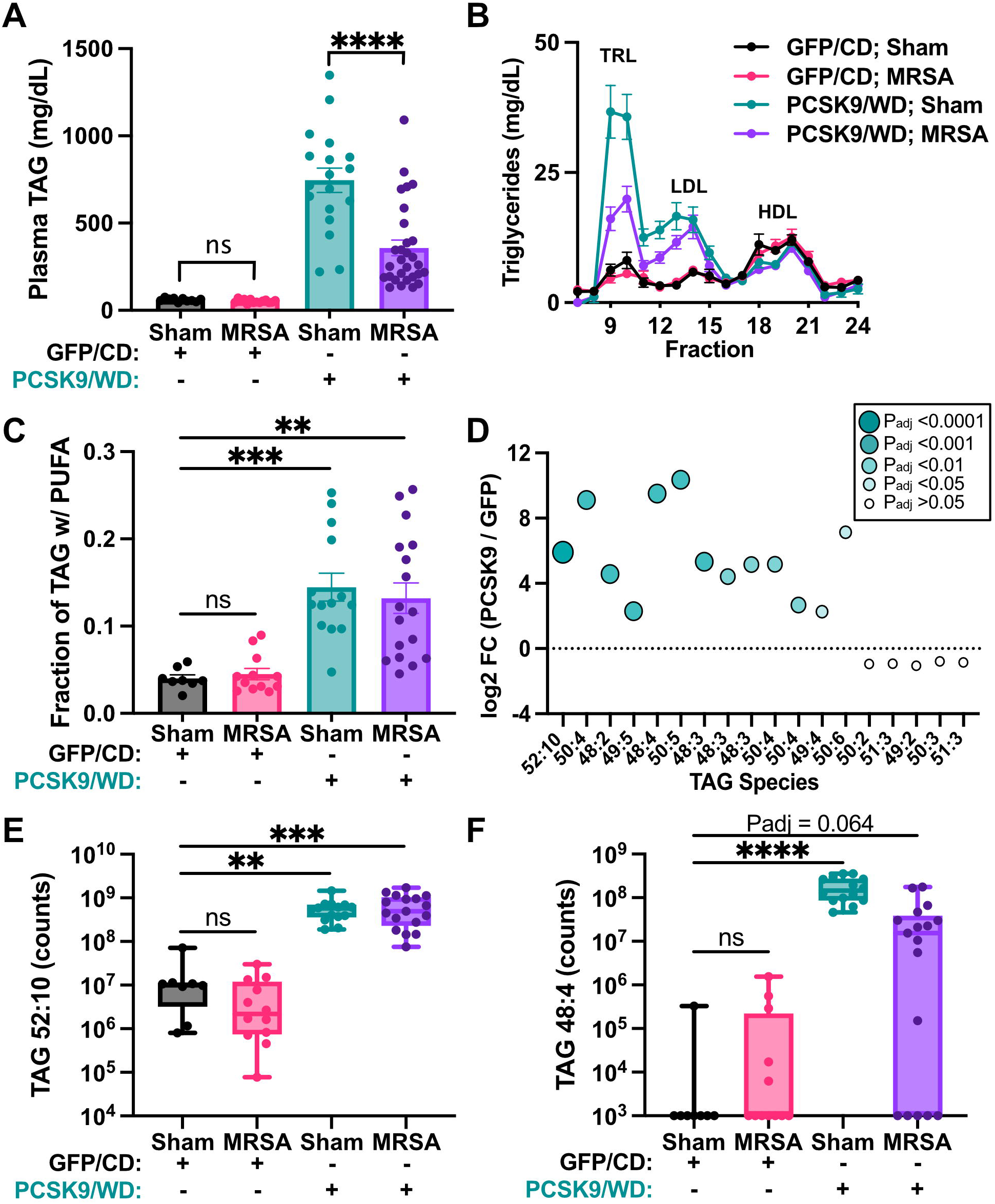
Western diet feeding increases circulating PUFA in plasma TAG. **A**) Plasma triacylglyerides (TAG) levels at harvest. AAV-GFP/CD+Sham, n=12; AAV-GFP/CD+MRSA, n=17; AAV-PCSK9/WD+Sham, n=19; AAV-PCSK9/WD+MRSA, n=29. Two-way ANOVA with Sidak’s post-hoc test, ****P_adj_ <0.0001. **B**) Plasma was fractionated by size-exclusion chromatography and TAG in each fraction were determined by colorimetric kit. **C**) Raw counts of TAG species containing at least one polyunsaturated fatty acyl chain were summed and divided by the sum of all TAG species. Note samples with insufficient volume or excessive hemolysis were not submitted for lipidomics analysis: AAV-GFP/CD+Sham, n=8; AAV-GFP/CD+MRSA, n=12; AAV-PCSK9/WD+Sham, n=14; AAV-PCSK9/WD+MRSA, n=17. Kruskal-Wallis test with Dunn’s multiple comparison test. **P_adj_ <0.01; ***P_adj_ <0.001. **D**) Relative enrichment of TAG species containing at least one polyunsaturated fatty acyl chain in AAV-PCSK9 mice fed WD relative to AAV-GFP mice fed CD. Student’s t-test (unpaired, two-tailed) with False Discovery Rate adjusted *P*-value (α<0.05). **E,F**) Raw counts of selected TAG species detected in each group. Note: Predicted fatty acyl chains of (**E**) 52:10 are 14:0_19:5_19:5; Predicted fatty acyl chains of 48:4 (**F**) are 12:0_16:0_20:4. Kruskal-Wallis test with Dunn’s multiple comparisons test. **P_adj_ <0.01; ***P_adj_ <0.001; ****P_adj_<0.0001. Data are mean +/- standard error of the mean (SEM).

### TRL, but not other host lipoproteins, are toxic to S. aureus in vitro

We next sought to determine whether host lipoproteins could contribute to pathogen toxicity *in vitro*. Prior studies have shown that exposure to host lipoproteins does not impede *S. aureus* growth *in vitro* and recent work has demonstrated that *S. aureus* can harvest fatty acids from cholesterol-rich lipoproteins to support pathogen growth, and even use cholesterol to detoxify their environment^36,41,42^. However, these studies have mostly focused on introduction of host lipoproteins to cultures pre-exponential growth. As our *in vivo* model represents chronic *S. aureus* infection, we sought to evaluate the impact of host lipoproteins to pathogen viability after exponential growth (**Figure 3A**). Physiologically relevant concentrations of HDL and LDL added to 24h cultures did not impact pathogen viability as assessed by CFU (**Figure 3B**). When post-exponential cultures were supplemented with human TRL, though, we observed a dose-dependent decrease in CFU (**Figure 3C**). Of critical importance, the experimental dose of 100-200 μg/mL of TRL protein represents physiologically normal to borderline-high range of fasting TRL in humans. As work from independent groups have reported no effect of TRL on *S. aureus* growth when added before exponential growth^42^, we modified our experimental design to compare matched addition of TRL at the initial dilution or 24h later. TRL supplementation at initial dilution modestly reduced CFU after 24h, but 48-72h cultures were comparable to vehicle (**Figure 3D**). Conversely, matched samples incubated with TRL after exponential growth resulted in a nearly 10-fold reduction in CFU at 48h and 72h time-points (**Figure 3D**). Thus, TRL-induced toxicity appears to be dependent on a pathogen-derived factor that accumulates in post-exponential cultures.

**Figure 3.**
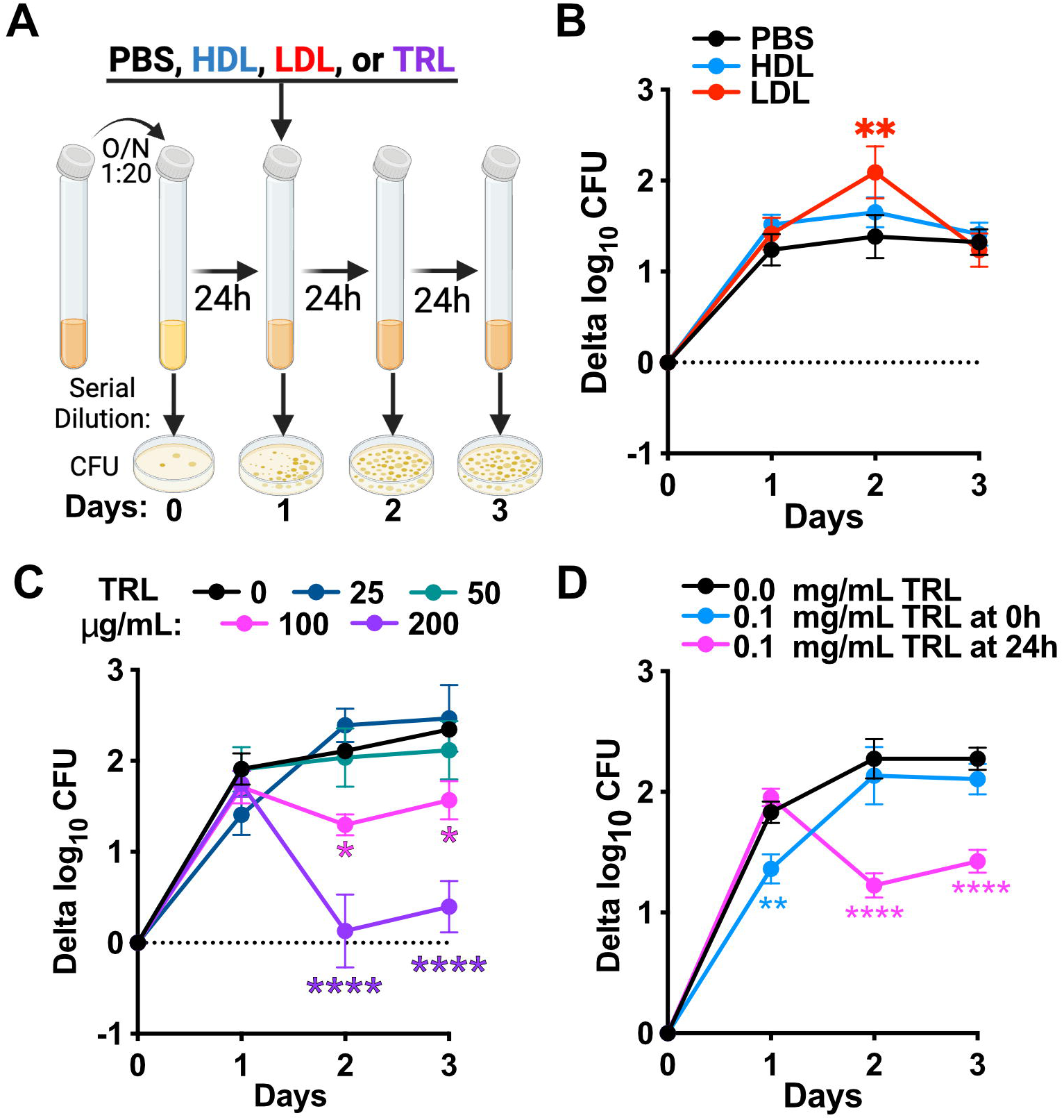
TRL, but not other host lipoproteins, are toxic to *S. aureus in vitro.* **A**) An overnight culture of CA-MRSA USA300 isolate LAC was diluted 1:20 in fresh TSB. Immediately after dilution, and every 24h thereafter, an aliquot was serially diluted and plated on TSA to determine CFU. **B)** Immediately after taking the aliquot on day 1, cultures were treated with phosphate buffered saline (PBS), 1.0 mg/mL HDL, or 0.5 mg/mL LDL for 48h (n=6 biological replicates/treatment). Two-way ANOVA with Dunnett’s multiple testing correction. **P_adj_<0.01, LDL vs PBS on day 2. **C)** Design as (A), but varying doses of TRL were added to 24h cultures on day 1 (n=5 biological replicates/treatment). Two-way ANOVA with Dunnett’s multiple testing correction. *P_adj_<0.05, 100 μg/mL TRL vs PBS; ****P_adj_<0.0001, 200 μg/mL TRL vs PBS. **D)** Design as (A), except that matched 100 μg/mL TRL were added at initial dilution (blue) or 24h culture (pink; n=6 biological replicates/treatment). Two-way ANOVA with Dunnett’s multiple testing correction. **P_adj_<0.01, 100 μg/mL TRL at 0h vs PBS; ****P_adj_<0.0001, 100 μg/mL TRL at 24h vs PBS.

### TRL exploit S. aureus secreted lipases to induce toxicity in vitro

Virulence factor production during the post-exponential phase is heavily influenced by the accessory gene regulator (*agr*) quorum-sensing system^52^. Secreted auto-inducing peptide (AIP; encoded by *agrD*) binds to the transmembrane receptor AgrC and triggers phosphorylation and activation of the transcription factor AgrA. Activated AgrA binds to promoter element P2 to amplify expression of the *agr* operon (*agrBDCA*) and to promoter element P3 to induce the non-coding RNA regulator and effector *RNAIII*, which promotes translation of a diverse arsenal of virulence factors (e.g., hemolysins, proteases, nucleases, lipases) and suppresses expression of adhesion proteins (e.g., staphylococcal protein A, fibronectin binding proteins). Previous studies have shown that host apolipoprotein B (ApoB), the structural scaffold of TRL and LDL, can disrupt *agr* quorum-sensing and virulence factor production through binding and sequestration of AIP^41–43^. It is plausible that hyperlipidemia-induced repression of *agr* and toxin production could contribute to reduced cortical bone destruction in AAV-PCSK9/WD mice (**Figure 1D,E**), but *agr* is dispensable for growth and Δ*agr* mutants of LAC have failed to influence CFU in chow-fed wild-type mice using our murine osteomyelitis model^53^. We therefore speculated that the *agr* regulon may induce a factor capable of interacting with, or modifying, host TRL.

We first validated that human TRL dose-dependently suppressed *agr* activity *in vitro* using LAC stably transformed with a P3-driven luciferase reporter. Growth, as determined by OD600, was minimally influenced by the presence of TRL (**Figure 4A**), whereas P3-driven luminescence at 24h was repressed in a dose-dependent manner (**Figure 4B**). Recent work from our group has shown that secreted lipases *lip* and *geh* (glycerol ester hydrolase) are among the most abundant proteins present in post-exponential conditioned media of LAC^54^, and expression of each lipase is amplified as part of the *agr* regulon through actions of *RNAIII*^55^. In cell lysates of stationary phase LAC treated with TRL, we observed dose-dependent reductions in mRNA expression of *geh* and *lip* that are consistent with disrupted *agr* signaling (**Figure 4C,D**). From these data, we rationalized that TRL-dependent suppression of lipases may benefit the pathogen by mitigating the release of toxic PUFA that are inert when stored as TAG. To test this idea, we titrated increasing concentrations of the covalent lipase inhibitor Orlistat into 24h cultures of LAC immediately prior to exposure to TRL (**Figure 4E**). Our results support that extracellular lipase activity is required for TRL-induced toxicity in LAC. It is notable that high doses of Orlistat were required for protection against TRL^23^, which likely reflects the high abundance of secreted lipases in post-exponential conditioned media of LAC.

**Figure 4.**
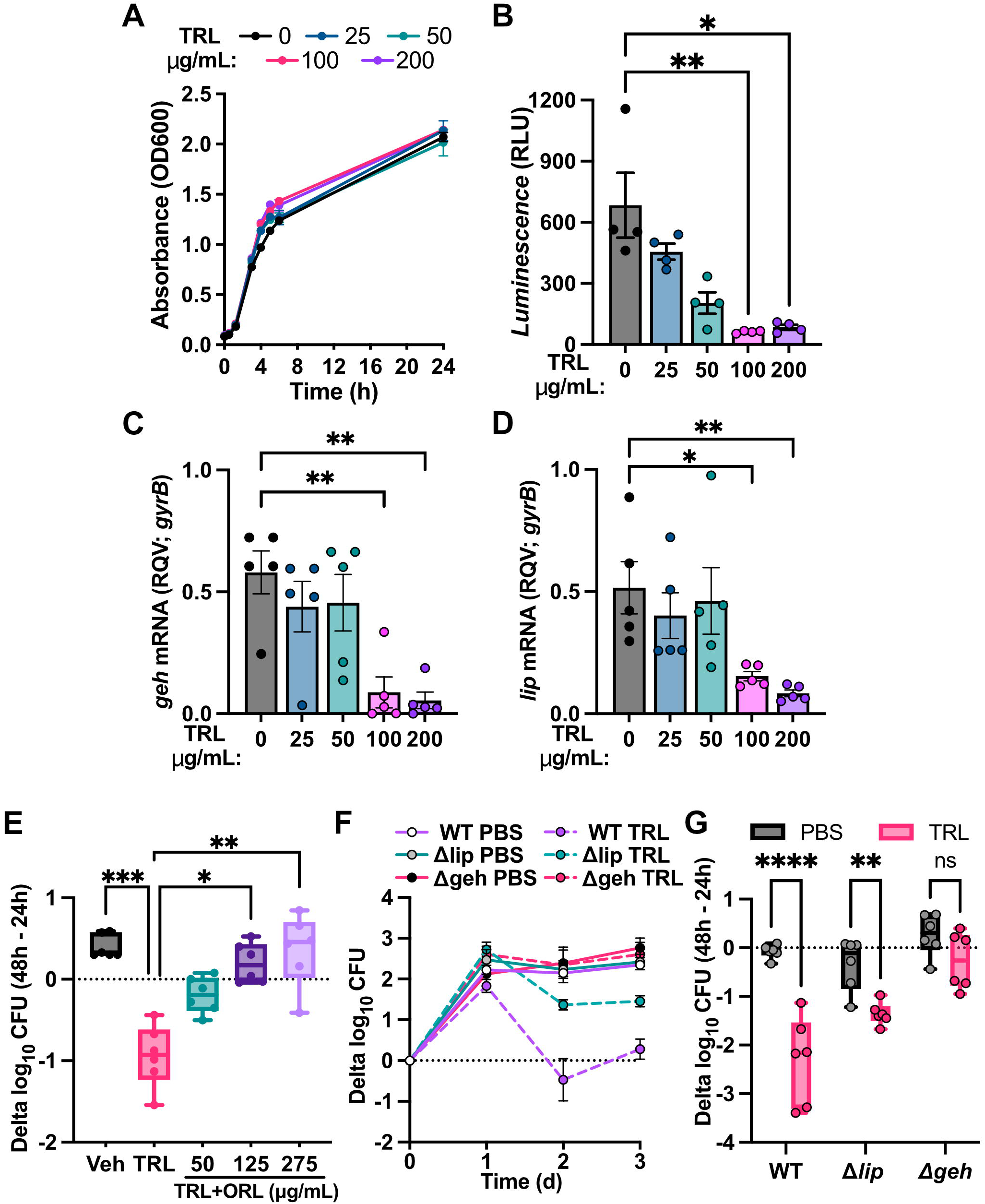
TRL exploit secreted lipases of *S. aureus* to induce toxicity *in vitro*. **A)** Overnight cultures of LAC stably transformed with a P3-driven *lux* reporter plasmid were diluted 1:30 into 14 mL round-bottom tubes containing TSB supplemented with PBS (vehicle) or the indicated concentration of TRL protein mass. OD600 was measured at baseline and various timepoints up to 24h (n=4 biological replicates). **B)** At 24h, P3-dependent luminescence values were recorded (n=4 biological replicates). Kruskal-Wallis test with Dunn’s multiple comparisons correction. *P_adj_<0.05, **P_adj_<0.01. **C-D)** Overnight cultures of LAC were diluted 1:20 into 14 mL round-bottom tubes containing TSB and cultured for 24h. These post-exponential cultures were then treated with PBS (vehicle) or indicated concentrations of TRL protein mass for 24h. Thereafter, cells were pelleted and total RNA extracted and mRNA expression of *geh* (**C**) and *lip* (**D**) was determined relative to *gyrB* and plotted as relative quantitative value (RQV). Kruskal-Wallis test with Dunn’s multiple comparisons correction. *P_adj_<0.05, **P_adj_<0.01. **E)** Design similar to (C-D) but indicated concentrations of the covalent lipase inhibitor Orlistat was added 15 minutes before addition of 200 μg/mL TRL at 24h (n=6 biological replicates/treatment). Data represent the change in CFU 24h after addition of TRL. Kruskal-Wallis test with Dunn’s multiple comparisons correction. *P_adj_<0.05, **P_adj_<0.01, ***P_adj_<0.001. **F)** Design as (B) with 200 μg/mL TRL added at 24h for cultures of wild-type (WT) LAC, Δ*lip* mutant LAC, or Δ*geh* mutant LAC (n=6 biological replicates/treatment). **G)** Change in CFU 24h after addition of TRL to WT LAC, Δ*lip* mutant LAC, or Δ*geh* mutant LAC cultures of (H). Two-way ANOVA with Sidak’s multiple comparison test. **P_adj_<0.01, ****P_adj_<0.0001.

Although both *lip* and *geh* are highly abundant in conditioned media of LAC, *lip* is reported to have specificity for short- and medium-chain fatty acids^33^, whereas *geh* exerts lipolytic activity on a broad range of substrates^21,23^. To determine their relative contributions to TRL-induced toxicity we repeated our experimental design in Δ*geh* and Δ*lip* mutant LAC (**Figure 4F**). Wild-type LAC and Δ*lip* mutants each showed significant reductions in CFU following post-exponential exposure to TRL, whereas Δ*geh* mutant LAC cultures were not altered for up to 48h post exposure to TRL (**Figure 4G**). Taken together, these data support that TRL-induced toxicity *in vitro* is dependent on lipolytic activity of Geh.

### Host lipase activity can contribute to TRL-induced toxicity

Disruption of *agr* quorum-sensing by ApoB has been proposed as a host adaptation to suppress virulence factor production during infection^41^. Yet, it could be interpreted from data presented here that AIP binding to ApoB is an elegant adaptation of the pathogen to monitor the presence of host-derived TAG substrates in the local environment and adjust lipase secretion to avoid toxicity. If this model of host response evasion were accurate, we hypothesized that host secretion of TRL-targeting lipase could be an effective counter measure in the infection microenvironment. Activated macrophages are known to express and secrete lipoprotein lipase (LPL) in the context of atherosclerosis^56,57^, but to our knowledge, there is limited data on LPL’s role in *S. aureus* infection.

To determine the potential impact of LPL to TRL-induced toxicity of *S. aureus*, we first diluted an overnight culture of bioluminescent LAC::*lux* in flasks containing TSB supplemented with PBS (vehicle), TRL, or TRL with recombinant lipoprotein lipase (rLPL). Flasks were used to provide a 10:1 air-to-liquid ratio that supports maximal *agr* activity^58–60^. Under these conditions the presence of TRL did not alter exponential growth as determined by OD600 with all cultures reaching post-exponential phase within 6h (**Figure 5A**). Endogenous luminescence from *lux* requires the reduced form of flavin adenine dinucleotide (FADH_2_) as a substrate and is therefore often used to evaluate the metabolic activity of *S. aureus*^61–64^. As expected, *lux* activity rose sharply during the exponential phase, reflecting strong expression of *lux* and availability of endogenous FADH_2_ at the 6h time point (**Figure 5B**). However, *lux* activity was repressed by TRL alone during and after the exponential phase, and this was further lowered by rLPL co-treatment where luminescence did not exceed background at 24h. Cultures that were serially diluted at 6h showed no difference in CFU between groups (**Figure 5C,D**), in agreement with OD600 values. Reduced luminescence at 24h in cultures treated with TRL alone were associated with a modest, but statistically significant, reduction in CFU, whereas toxicity was potentiated in cultures co-treated with TRL and rLPL (**Figure 5D**). Detection of ApoB in conditioned media of cultures supplemented with TRL decreased during post-exponential growth and were undetectable by western blot after 24h, which could be due to proteolytic degradation or oxidative damage to epitopes (**Figure 5E**). In wild-type LAC::*lux*, secretion of the *bona fide agr* target hemolysin-alpha (Hla; alpha-toxin) increased during exponential growth with high abundance of this toxin in conditioned media collected at 24h (**Figure 5E**). The presence of TRL alone strongly repressed Hla secretion to conditioned media during late exponential phase, consistent with abrogated *agr* signaling. Most intriguingly, though, TRL treatment did not appear to elevate secretion of extracellular staphylococcal protein A (eSpa), a key marker of biofilm formation that is commonly enhanced in Δ*agr* mutant strains^54,65–67^. These data further support that TRL exposure disrupts *agr-*mediated virulence factor secretion and introduces a novel observation that exogenous lipase activity can help create a toxic microenvironment for *S. aureus*.

**Figure 5.**
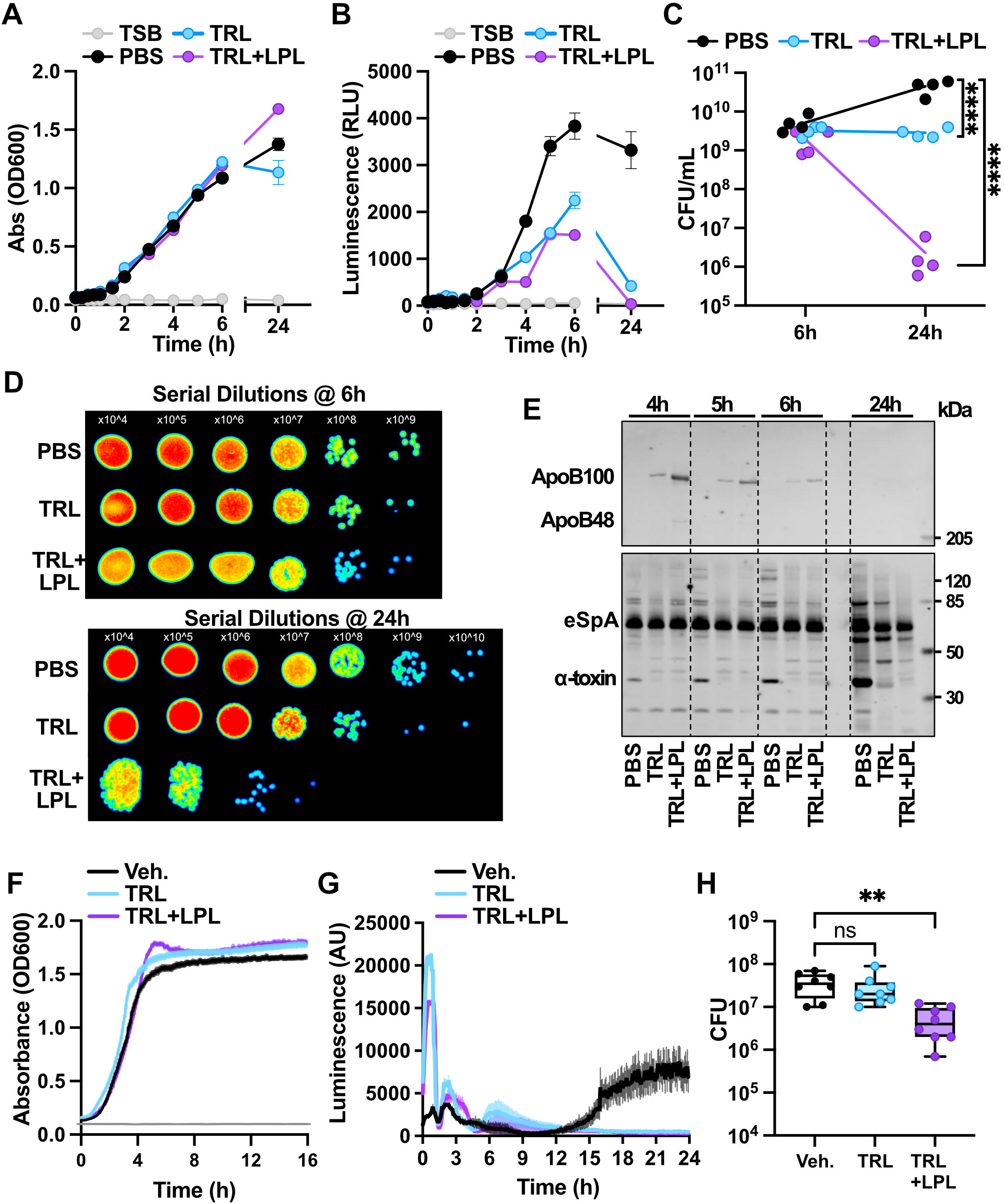
Host lipase activity can contribute to TRL-induced toxicity. **A-E)** *S. aureus* LAC::*lux* was diluted 1:30 into TSB in an Erlenmeyer flask supplemented with PBS or 200 μg/mL TRL with or without 2.5 μg/mL recombinant lipoprotein lipase (rLPL) as to maintain a 10:1 air-to-liquid ratio. **A)** Bacterial growth as determined by absorbance at OD600 (n=4 biological replicates). **B)** Endogenous bioluminescence reflecting cellular levels of *lux* and substrate FADH_2_ (n=4 biological replicates) **C)** Observed CFU/mL at 6 and 24h (n=4 biological replicates). Two-way ANOVA with Dunnett’s multiple testing correction, ****P_adj_ <0.0001. **D)** Relative *lux* activity of serially diluted cultures taken at 6h or 24h from (C), serially diluted and spotted overnight on TSA plates. **E)** Immunoblot of conditioned media collected at different time points of (A,B). (Top) Anti-apolipoprotein-B with reactivity to human ApoB100 and ApoB48 isoforms. (Bottom) Specific bands of anti-alpha hemolysin (*hla*; alpha-toxin) in conditioned media and non-specific bands of several proteins, including extracellular Staphylococcal protein A (eSpA). **F-H)** *S. aureus* LAC::*lux* was diluted 1:30 into TSB in a 96-well plate supplemented with PBS or 200 μg/mL TRL with or without 2.5 μg/mL recombinant lipoprotein lipase (rLPL) to maintain a nearly 1:1 air-to-liquid ratio. **F)** Bacterial growth as determined by absorbance at OD600 (n=6 biological replicates). **G)** Endogenous bioluminescence reflecting cellular levels of *lux* and substrate FADH_2_ (n=6 biological replicates). **H)** Observed CFU at 24h (n=6 biological replicates). One-way ANOVA with Dunnett’s multiple testing correction, **P_adj_ <0.01.

We considered that increased aeration in flask-grown cultures (**Figure 5A-E**; 10:1 air-to-liquid ratio) relative to round bottom tubes (3:1 air-to-liquid ratio; **Figure 3D**) contributed to increased sensitivity to TRL. Therefore, we next diluted an overnight culture of LAC::*lux* in a 96-well plate with a nearly 1:1 air-to-liquid ratio and continuously monitored either growth (**Figure 5F**) or luminescence (**Figure 5G**) in the presence of PBS, TRL or TRL with rLPL. Similar to cultures grown in flasks, TRL failed to augment exponential growth of LAC::*lux* as determined by OD600 in the 96-well plate format (**Figure 5F**). Activity of *lux*, though, was dramatically altered by the presence of TRL with multiple surges in *lux* activity during lag and exponential phases (**Figure 5G**). We postulate that increased luminescence at these early time points could be the result of recently described fatty acid dehydrogenase enzymes that generate FADH_2_ as they shorten exogenous long-chain fatty acids for incorporation to membrane lipids^18,19^. Luminescence declined to similar levels in all groups as cells transitioned out of exponential growth, but only cells treated with PBS recovered *lux* activity beyond post-exponential phase. As with cultures grown in flasks, luminescence in cells treated with both TRL and rLPL did not exceed background luminescence after 24h of culture. When plated for CFU, though, cultures treated with TRL alone were not significantly different from controls, whereas TRL with rLPL resulted in a >10-fold reduction in CFU (**Figure 5H**). We thus conclude that lipolysis of TRL by host lipases represents a novel, active mechanism of innate immunity against *S. aureus* that may have untapped therapeutic potential for chronic infection.

## DISCUSSION

Here, we show that host TRL actively promote toxicity to *S. aureus* that is dependent on extracellular lipase activity (**Figure 6**). Host lipoproteins, as well as the passive fatty acid carrier albumin, have been shown to permit or support pathogen growth in blood, and our data are largely supportive of these conclusions^36,41,42^. Our data are also consistent with previous observations that TRL passively contribute to host defense through suppression of virulence factor production through *agr*. However, novel data presented here introduce a paradigm shift in the function of TRL during infection, as we demonstrate that TRL have bactericidal properties that are engaged by lipase activity of the pathogen or the host. As *S. aureus* osteomyelitis often presents as chronic infections with periods of quiescence and flares, targeted manipulation of host TRL secretion and lipase activity may represent novel therapeutic strategies for this debilitating infectious disease.

**Figure 6.**
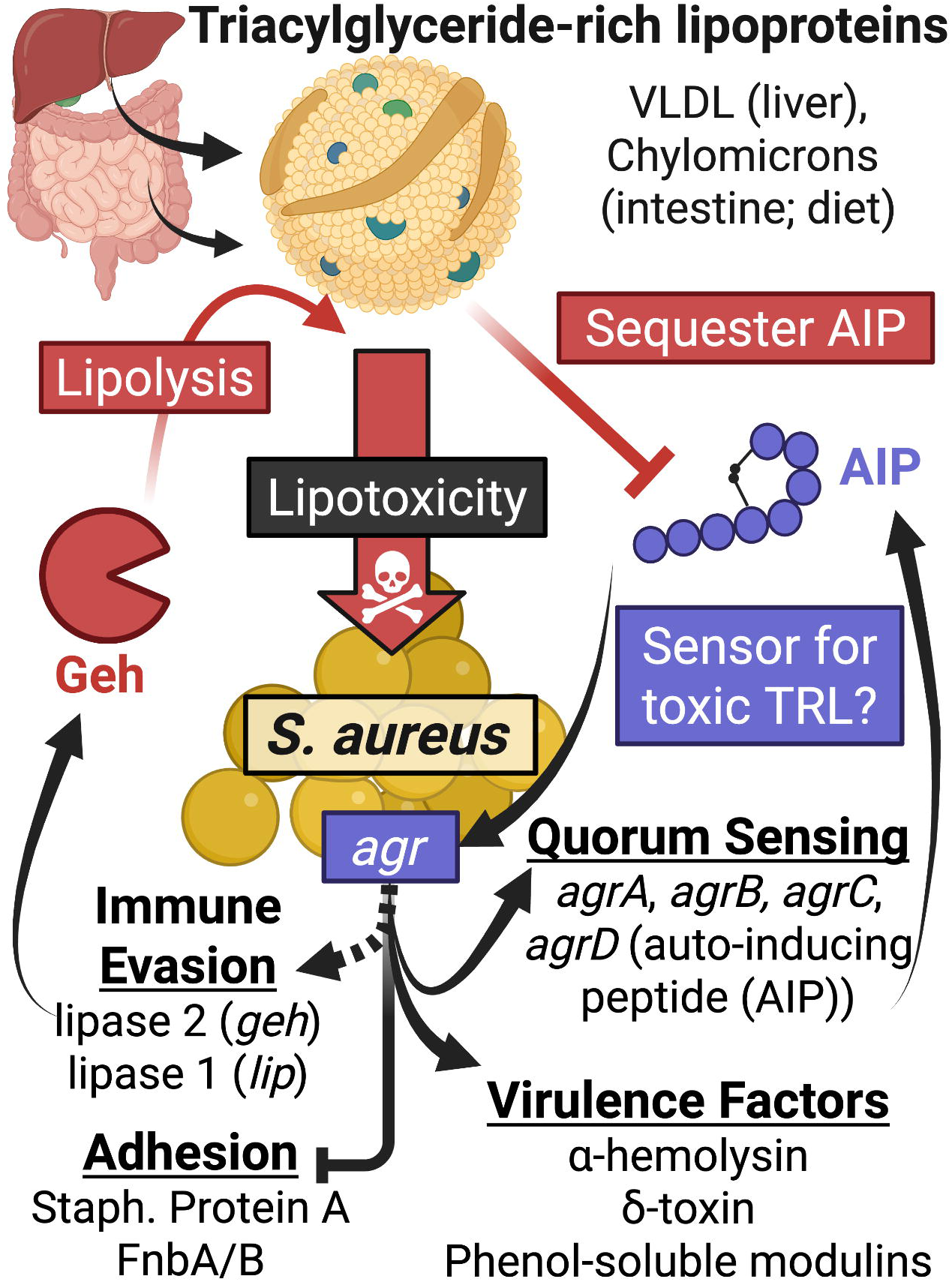
Host triacylglyceride-rich lipoproteins passively control *S. aureus* phenotype and actively contribute to lipotoxic cell death. Triacylglyceride-rich lipoproteins (TRL) are secreted to blood from the liver (VLDL) and intestine (chylomicrons) for systemic distribution. Previous studies support that apolipoprotein-B isoforms of TRL bind and sequester auto-inducing peptide (AIP) to disrupt accessory gene regulator (*agr*) quorum-sensing and virulence factor secretion, providing a passive form of host defense. New data presented herein support that host production of TRL actively contribute to host defense against *S. aureus* through lipotoxic stress that is dependent upon activity of extracellular lipases. These data suggest that manipulation of host lipoprotein metabolism and/or lipase activity may have untapped therapeutic potential for treatment of chronic *S. aureus* infections.

Our observation that increasing PUFA content in TRL are associated with reduced bacterial burden is consistent with the notion that *S. aureus* poorly tolerates PUFA-rich environments^28,29,31,32,68^. It should be noted, though, that milkfat source in the WD used is relatively poor in PUFA and is instead enriched in saturated fatty acids that could support *S. aureus* growth. Future studies should evaluate whether consumption of PUFA-rich food oils – and their systemic delivery by TRLs – further enhances the host response to chronic *S. aureus* infections. Reduced bacterial burden in the face of severe hypercholesterolemia in our mice is also intriguing, as recent work has suggested that Geh detoxifies PUFA-rich environments by esterifying toxic fatty acids to cholesterol^69^. Thus, future studies should also determine whether pharmacological inhibition of cholesterol absorption or abstaining from dietary cholesterol enhances antibacterial properties of hypertriglyceridemia.

Lipase activity is often enhanced in *S. aureus* isolates from deep tissue infection relative to superficial tissues^33^, and the genetic integrity of *geh* is intact in approximately 85% of sequenced strains^70,71^. It is possible that *geh^+^* strains are best adapted for deep tissue infections because of limited exposure to TRL and extracellular lipids of the host. Our observation that TRL alone seemed to repress *lux* activity – and by extension metabolic function - *in vitro* may suggest an evolved rationale for sustaining plasma TAG long after resolution of infection^40,72,73^. We believe our observed phenotype is consistent with suppressed *agr* quorum-sensing, and this may explain how weak *agr* signaling better supports vegetation formation in lipid-rich environments during endocarditis^74^. Future studies should investigate the influence of TRL to other infection sites and with varied clonal complexes.

Our finding that host lipase activity can compensate for TRL-mediated suppression of *agr* opens an exciting new possibility for active modulation of the host response. Fibrates are synthetic agonists of peroxisome-proliferator activated receptor alpha (PPARα) and a first-line therapy for hypertriglyceridemic patients^75,76^. Through activation of PPARα, fibrates induce hepatic secretion of TRL and systemic expression of *LPL* to promote lipolysis of circulating TRL and movement of hydrolyzed fatty acids into tissues for β-oxidation. We thus speculate that fibrate use in patients with chronic *S. aureus* infection may both reduce cardiovascular risk by lowering TRL in circulation and promote a lipid-rich extracellular environment within infected tissues. Unfortunately, the effects of TRL are likely dependent on vascular supply to infected tissues, the lack of which is a common cause of chronic *S. aureus* infection in patients with type II diabetes^77–79^. Nevertheless, it is intriguing that recent studies have shown that pharmacologic activation of PPARγ hastens resolution of *S. aureus* skin infections in mice by promoting M2-like macrophage polarization^80,81^. Similar to PPARα, PPARγ activation promotes LPL secretion from macrophages^82,83^, which can provide fatty acid substrates necessary for oxidative metabolism in M2-like macrophages. We therefore hypothesize that PPARγ-driven expression of *Lpl* could directly contribute to enhanced wound recovery in skin infections, but this remains to be determined.

## RESOURCE AVAILABILITY

### Lead Contact

Further information and requests for resources and reagents should be directed to, and will be fulfilled by, the lead contact, Ryan M. Allen.

### Materials availability

Materials generated in this study are available upon request from the lead contact in accordance with existing guidelines and regulations.

### Data and code availability

All data supporting the findings of this study are available within the article and its supplemental information. Any additional information required to reanalyze the data reported in this work paper is available from the lead contact upon request.

## ACKNOWLEDGEMENTS

The authors would like to thank Kirsten LeCompte, Ahmed Tolba, Rodrigo Meade, and Drs. Mara Campbell, Sanja Novak, Amy Sato and Neha Dole for technical assistance and helpful discussions. We also thank faculty of the University of Arkansas for Medical Studies (UAMS) Center for Microbial Pathogenesis and Host Inflammatory Responses for consultation on experimental design and interpretation. This work is supported by a UAMS Honors in Research fellowship (C.N.C.), UAMS Translational Research Innovations and Partners Fellowship (C.N.C), UAMS Sturgis Foundation Award (R.M.A), Pilot Funding from the Center for Microbial Pathogenesis and Host Inflammatory Responses (R.M.A) supported by the National Institutes of General Medical Sciences Center of Biomedical Excellence (COBRE) grant 5P30GM145393 (M.S.S), National Institutes of Allergy and Infectious Disease Grant R01AI119380 (M.S.S.), and Department of Veterans Affairs Grant BX006469 (A.J.M.).

## AUTHOR CONTRIBUTIONS

Conceptualization: R.M.A., M.S.S., C.N.C.; Methodology: R.M.A., A.J.M., M.S.S., K.E.B.; Investigation: C.N.C., T.V., A.N., A.H., N.J.T., R.M.A.; Writing – original draft: T.V., C.N.C., R.M.A.; Writing – reviewing and editing: T.V., C.N.C., K.E.B., M.S.S., N.J.T., A.J.M., A.H., A.N., R.M.A.; Visualization: C.N.C., T.V., R.M.A.; Supervision: A.J.M., K.E.B., M.S.S., R.M.A.; Funding Acquisition: C.N.C., R.M.A., M.S.S.

## Supplemental information

Figure S1-S3

Supplemental Table S1

Supplemental Table S2

## STAR METHODS

### Resource Table

| Reagent or Resource | Source | Identifier |
| --- | --- | --- |
| **Recombinant proteins, antibodies, AAV vectors, plasmids, and chemicals** |  |  |
| Rabbit anti-Apolipoprotein-B | Proteintech Group | 20578-1-AP |
| Rabbit anti-Apolipoprotein-A1 | Proteintech Group | 14427-1-AP |
| Rabbit anti-Apolipoprotein-E | Proteintech Group | 18254-1-AP |
| Rabbit anti-hemolysin alpha | Sigma-Aldrich | S7531 |
| Anti-Rabbit IgG Dylight 680 | Cell Signaling Technologies | 5366S |
| Anti-Mouse IgG Dylight 800 | Cell Signaling Technologies | 5257S |
| Orlistat | Sigma-Aldrich | O4139 |
| AAV8-GFP | VectorBuilder | VB010000-9287ffw |
| AAV8-PCSK9-D377Y | LSU Health Sciences<br>Center - Shreveport | RRID:SCR_027498 |
| Lysostaphin | MedChemExpress | HY-92329 |
| Oil Red O | Sigma-Aldrich | O0625 |
| Lipoprotein Lipase | R&D Systems | 9888-LL |
| pMk4-luxABCDE::agrP3 | This study |  |
| **Critical Commercial Assays and Kits** |  |  |
| Cholesterol Reagent | Pointe Scientific | C7510 |
| Triglycerides Reagent | Pointe Scientific | T7532 |
| SYTO RNASelect Green | Invitrogen | S32703 |
| Hematoxylin and Eosin Kit | Abcam | ab245880 |
| BCA Protein Assay Kit | Thermo Scientific Pierce | 23227 |
| **Experimental models: plasma** |  |  |
| Human Recovered Plasma,<br>Frozen, K<sub>2</sub>-EDTA | ZenBio | SER-PLE |
| **Experimental models: strains and organisms** |  |  |
| C57BL/6J mice | Jackson Laboratory | #000664 |
| *S. aureus* USA300 LAC | Mark Smeltzer | Wormann et al. 2011 |
| *S. aureus* USA300 JE2 Δgeh | Nebraska Transposon<br>Mutant Library | NE1775 |
| *S. aureus* USA300 JE2 Δlip | Nebraska Transposon<br>Mutant Library | NE338 |
| *S. aureus* USA300 LAC::*lux* | This study |  |
| *S. aureus* USA300 LAC Δ*geh* | This study | |
| *S. aureus* USA300 LAC Δ*lip* | This study | |
| **Oligonucleotides** |  |  |
| Primers for cloning and qPCR,<br>see Table S2 | Integrated DNA<br>Technologies | N/A |
| **Software** |  |  |
| Fiji (ImageJ) Version 2.14 | Schindelin et al. 2012 | https://imagej.net/software/fiji |
| Prism 10 Software | GraphPad | https://www.graphpad.com |
| Skyscan CT-Analyzer | Bruker | https://bruker.com |
| Skyscan N-Recon | Bruker | https://bruker.com |
| Biorender | Biorender | http://www.biorender.com |

### Experimental models

#### *S. aureus* USA300 and generation of *S. aureus* mutants

Community-acquired methicillin-resistant *Staphylococcus aureus* (CA-MRSA) isolate USA300 originally isolated from the Los Angeles County jail (LAC) was used for most experiments. Bioluminescent LAC::*lux* was previously generated by phage-mediated transduction of an integrated plasmid pRP1195 from USA300 strain NRS384, a generous gift from Dr. Roger Plaut^63^. Transposon mutants for *geh* and *lip* in USA300 JE2 were acquired from the Nebraska Transposon Mutant Library (NTML) and transferred to LAC using φ11-mediated transduction^84^. Antibiotics used in selection of different strains and transposon mutants: chloramphenicol 10 μg/mL (LAC::*lux*); lincomycin 25 μg/mL; erythromycin 5 μg/mL. Validation of transposon insertion was determined by PCR using gene-specific primers (Supplemental Table S2). For all experiments, LAC, LAC::*lux* and LAC mutants were cultivated in tryptic soy broth (TSB) at 37°C with shaking at 200 rpm and plated on tryptic soy agar (TSA) without antibiotic. For most *in vitro* experiments, 3-3.5 mL of culture was grown in 14.2 mL round bottom polypropylene tubes with a loosely attached lid. Where indicated, 13 mL of culture was grown in 125 mL glass Erlenmeyer flasks with a foil cap. Alternatively, 200 μL of culture was grown in white 96-well plates with a flat, optically clear bottom and sealed with Breathe-Easy film (USA Scientific). Absorbance at OD600 and endogenous bioluminescence were measured using a Clariostar plate reader (BMG LabTech).

For *in vivo* experiments, LAC was grown overnight at 37°C in TSB with shaking at 200 rpm and pelleted the following morning. The pellet was washed three times with sterile phosphate buffered saline (PBS) and then resuspended in PBS at a density of 5 x 10^8^ colony forming units (CFU) per mL. The concentration was confirmed by plating serial dilutions on TSA. For infection, 2 μL of this suspension (1 x10^6^ CFU) was used per mouse.

### Ethics statement

Animal protocols were approved by and performed according to the regulations of the University of Arkansas for Medical Sciences Institutional Animal Care and Usage Committee (IPROTO202200000450, IPROTO202200000484). All animal studies were conducted in accordance with NIH guidelines, the Animal Welfare Act, and United States federal law.

### Animal models

Male and female C57BL/6 mice, aged 7-10 weeks, received either a control adeno-associated viral vector (AAV8-GFP, 1×10^11^; synthesized by VectorBuilder) or an AAV to promote over-expression of gain-of-function mutant (D377Y) of proprotein convertase subtilisin/kexin type 9 (AAV8-PCSK9, 1×10^11^) ^45^. AAV8-PCSK9 was produced using the Addgene pAAV/D377Y-mPSCK9 vector (#58376) by the AAV Virus Production Core (RRID:SCR_027498) in the Center for Cardiovascular Diseases and Sciences at LSU Health Shreveport. Mice receiving AAV8-GFP (n=29; sham=12, MRSA=17) were then fed a chow diet for 14 weeks, whereas AAV8-PCSK9 mice (n=47; sham=18, MRSA=29) were fed a western diet containing 42% fat and 0.2% cholesterol (TD.88137, Envigo). At 14 weeks, mice in each cage received a traumatic injury to the femur, as previously described^46^. First, mice were anesthetized with isoflurane and a surgical incision was made to expose the flat surface of the femur where a unicortical defect was bore with an 18G needle. The hole was then inoculated with 2 μL containing 10^6 CFU of USA300 LAC strain of Methicillin-resistant *Staphylococcus aureus* (prepared as described above), or 2 μL of sterile saline (Sham). Muscle and skin were sutured, and animals were immediately returned to their respective cages. To mitigate bias, each cage contained a mixture of mice receiving sham or MRSA. Body weight was measured before surgery and 3 days post-surgery for all mice and there were no observed complications. All mice were humanely euthanized at 14d post-surgery for collection of blood and tissues. Femurs were wrapped in sterile-saline soaked gauze and stored at −80°C prior to microcomputed tomography. After imaging, femurs were homogenized and bacterial burdens were determined as described below.

## METHOD DETAILS

### Analysis of plasma lipids and lipidomics

Blood was collected immediately after euthanasia by cannulation of the vena cava in a syringe containing EDTA. Whole blood was stored on ice prior to centrifugation at 2,500 x *g* for 10 minutes at 4°C. The plasma supernatant was transferred to a separate tube and used fresh for analysis or stored at −80°C for down-stream processing. Plasma was assayed for total cholesterol and triacylglycerides using colorimetric kits and quantified according to a standard curve (Pointe Scientific). For assessment of lipid/lipoprotein distribution, plasma samples (100 μL) were diluted in sterile saline and filtered on a 0.22 micron column prior to injection to an AKTA Pure system containing two tandem Superose-6 gel filtration columns (Cytiva). A total of forty 1.5 mL fractions were collected, with the first seven fractions representing the void volume and albumin clearance by fraction 23. For each fraction (7-24) 75 μL was assayed directly for total cholesterol and triacylglycerides using colorimetric kits (Pointe Scientific). Values for each fraction were calculated using a standard curve. For the detection of RNA, 50 μL of each fraction was incubated with SYTO RNASelect (Invitrogen) at 37°C for 30 minutes prior to detection of fluorescence on a plate reader, as previously described^85^.

Lipidomics studies were conducted using 50 μL of plasma in samples with low hemolysis. Sample extraction, data processing and reporting were done as described recently with some modifications^86^. In brief, untargeted lipidomics was performed using a Vanquish Horizon UHPLC system with an Accucore C30 column (2.1 × 150 mm, 2.6 µm) at 45 °C, employing mobile phases of ACN/H₂O (60:40) and IPA/ACN (90:10), each containing 10 mM ammonium formate and 0.1% formic acid, at a flow rate of 260 µL/min and an 8 µL injection volume. Samples were maintained at 4 °C, and separation was achieved using a gradient from 32% to 97% B over 35 min. Mass spectrometry was conducted on an Orbitrap Exploris 480 with HESI in separate positive and negative modes (spray voltage 3.6/3.0 kV; vaporizer 350 °C; ion transfer 320 °C), acquiring full MS scans at 120,000 resolution (m/z 100–1500) followed by Top-5 DDA MS/MS using a 1.0 m/z isolation window and stepped HCD energies of 20/30/40%. MS/MS spectra were collected at 15,000 resolution with dynamic exclusion and isotope filtering, and routine QC injections ensured instrument stability. Raw data were processed in MS-DIAL v4.9.221218 using standard tolerances, MSP library matching, and in-silico MS/MS annotations, with features removed if sample-to-blank <10 or QC RSD >30%. Only lipids detected in >80% of QC samples and supported by MS/MS confirmation were retained for final statistical analysis. Data were collected by running the samples separately in positive and negative ionization modes. Triacylglycerides were identified as their ammoniated adducts from the positive mode datasets.

### Isolation of lipoproteins by density-gradient ultracentrifugation

Frozen human recovered plasma were purchased from Zenbio and stored at −80°C. Individual samples were thawed at room temperature and fibrous aggregates were removed by vacuum filtration through a 0.45-micron filter. Sequential density gradient ultracentrifugation (DGUC) was performed using a Beckman Coulter Optima XPN-80 Ultracentrifuge with an SW32Ti swinging bucket rotor. Filtered plasma density was adjusted with potassium bromide to float TRL (<1.019 g/mL), LDL (1.019-1.063 g/mL) and HDL (1.064-1.022 g/mL) in sequential runs, as described previously^87^. Isolated lipoproteins were immediately transferred to Snakeskin dialysis tubing (3.5 kDa cutoff; ThermoScientific) and dialyzed in 1x PBS for at least four changes of 200-fold excess buffer. After dialysis, LDL and HDL were concentrated using Amicon Ultra filtration units (10 kDa molecular weight cut-off; Millipore) and then immediately sterile filtered (0.22 micron) by syringe. Concentration by ultra-filtration was necessary for some, but not all TRL samples. Total protein concentration was determined by standard curve using the BCA Protein Assay Kit (Pierce). Where indicated, triacylglycerides were assessed by colorimetric kit (Pointe Scientific).

### Assessment of atherosclerosis

After euthanasia with isoflurane and blood collection via cannulation of the vena cava, mice were perfused with PBS until the vasculature appeared clear. The heart, including the attached aortic root, was harvested, embedded in optimal cutting temperature (OCT) compound, and snap-frozen on dry ice^88^. Following guidelines from Paigen et al., fifty 7-μm sections were collected from the onset of the aortic root—identified by the first appearance of the valve cusps—using a Microm HM550 cryostat. Oil Red O (ORO) staining was applied to two sets of paired sequential tissue sections spaced 70 μm apart, beginning approximately 150 microns from the first appearance of the valve cusps^47^. Adjacent sections were processed for hematoxylin and eosin (H&E) staining (Abcam). Stained sections were imaged using a Nikon Eclipse microscope and analyzed using Fiji v2.14.0 to calculate total lesion area. Within these lesions, necrosis was quantified as a percentage of the total lesion area.

### MicroCT assessment of bone disease and quantification of bacterial burden

Frozen femurs were scanned using Skyscan 1275 X-ray Microtomograph (Bruker, Kontich, Belgium) and analyzed as previously described^89^. Reconstruction was done using the Skyscan N-Recon software. The Skyscan CT-analyzer software was used to process the reconstructed cross-sectional slices and create preliminary ROIs of just the cortical bone using a semiautomated protocol, as follows: global thresholding (lowL=L90; highL=L255), round closing in 3D space pixel size 4, round opening in 3D space, pixel size 1, round closing in 3D space, pixel size 8, and round dilation in 3D space pixel size 3. To keep the cortical bone, we drew inclusive or exclusive contours on the periosteal surface that allowed for the correction of the resultant images, which were loaded as ROI. The amount of cortical bone destruction will be evaluated by deducting the value obtained from each bone from the average obtained from sham operated bones infected with sterile PBS. The volume of cortical bone will be determined using these defined ROIs, with a threshold of 70–255. New bone formation was quantified using the subtractive ROI function on the previously delineated cortical bone ROI images and calculating the bone volume included in the newly defined ROI using a threshold of 45–135.

Each femur sample was homogenized with a Bullet Blender 5 Gold (Next Advance Inc., Troy, NY) and reconstituted in 2 ml of sterile PBS immediately following μCT imaging. An aliquot of this suspension was serially diluted in sterile PBS and each dilution was spotted on TSA without antibiotic selection. Plates were incubated at 37°C overnight and CFUs were manually counted the following day.

### RNA isolation from tissue and S. aureus cultures

Frozen liver samples were homogenized in TriReagent (Zymo) with cubic zirconium beads and an oscillating bead beater (Qiagen). A 0.2 volume of chloroform per 1 volume TriReagent was added to the homogenate and mixed by inversion for 2 minutes at room temperature. Samples were spun in a centrifuge at 12,000 x *g* for 15 minutes and the aqueous phase was transferred to a new tube and mixed with 1.5 volumes of 100% ethanol by vigorous shaking. Samples were then incubated on ice for 5 minutes and then applied to an RNA Clean and Concentrate column (Zymo). Washing and elution steps were completed according to manufacturer instructions. To isolate RNA from *S. aureus*, cultures were first pelleted by centrifugation, conditioned media removed by pipette. Cell pellets were then digested with lysostaphin (10 μg) at 37°C for 30 minutes prior to transfer to ice and addition of 700 μL of TriReagent. Digested cells were then pulverized at 4°C with cubic zirconium beads in a bead beater (2 cycles of 20 seconds; Qiagen). Chloroform was added as described above for phase separation, and RNA was isolated using the RNA Clean and Concentrate Kit as described above.

*Real-time PCR*. Total RNA from liver or bacterial cultures were quantified by NanoDrop and equal amounts of RNA from each sample were used for cDNA synthesis using the High-Capacity cDNA Synthesis Kit (Applied Biosystems). Quantitative real-time PCR was carried out using diluted cDNA and 2X Power SYBR Green master mix (Applied Biosystems) on a QuantStudio 7 Flex Real-Time PCR system (Applied Biosystems). Gene specific primers were designed and synthesized using PrimerQuest (Integrative DNA Technologies; Supplemental Table S2). Raw Ct values were used to calculate relative expression from reference genes (mouse = *Rplp01* (36b4); *S. aureus* = *gyrB*). Data are reported as relative quantitative values (RQV) calculated from the delta-Ct method.

### Immunoblot analysis

Individual fractions from size-exclusion chromatography (SEC) were mixed 5:1 with 5X RIPA buffer (ThermoFisher) to extract lipids from apolipoproteins, and then samples were denatured in Protein Sample Loading Buffer (LI-COR) supplemented with beta-mercaptoethanol at 70C for 5 minutes and then immediately placed on ice. Conditioned media samples were treated similarly, except these samples were not diluted in RIPA. Denatured protein samples were then run on NuPAGE 4-12% Bis-Tris or 3-8% Tris-acetate gels (Life Technologies) at 150V on ice with PageRuler Plus Pre-stained protein standards (Thermo Scientific) as molecular weight references. After electrophoresis, gels were transferred to PVDF membranes using the iBlot 3 system (Life Technologies). Membranes were then blocked in Fluorescence Blocking Buffer (Thermo Scientific) at room temperature for 30 minutes before dilution of the primary antibody. Primary antibodies were incubated for at least 1 hour at room temperature or overnight at 4C, washed three times in PBS supplemented with Tween-20 (0.05%) for 5 minutes. Similar procedures were repeated for secondary antibodies. Blots were imaged using the Odyssey Infrared Imaging System (LI-COR) and analyzed using Image Studio Lite and Image J software suites.

### Quantification and statistical analysis

Statistical method and sample size (n) for experiments are indicated in the corresponding figure legend. For all *in vitro* studies, n indicates the number of biological samples obtained from at least three independent experiments. For *in vivo* studies, n indicates the number of mice per group. Treatments for animal studies were randomized within individual cages to limit cage bias. For atherosclerosis studies, quantification of lesion area was assessed in a blinded fashion by at least two researchers. Statistical analysis was performed using Prism 10 Software (GraphPad) on the indicated biological replicates. Where two treatments were compared with a sample size >3 for each treatment a Mann-Whitney U-test (two-sided) was performed with α = 0.05. Where three or more groups were compared the non-parametric Kruskal-Wallis test with Dunn’s correction for multiple comparisons was used with α = 0.05. Where more than one test was performed for three or more groups a two-way ANOVA was performed using Sidak’s post-hoc test to control the False Discovery Rate (FDR; *q* = 0.05), or when multiple comparisons were made to a single group, Dunnett’s test was performed. Statistical significance was defined as p<0.05. Data are presented as mean +/- standard error of the mean (SEM).

