## Supplementary figures and images for "Lipolysis of host triacylglyceride-rich lipoproteins creates a toxic microenvironment for *Staphylococcus aureus*"

### Supplemental Figure S1

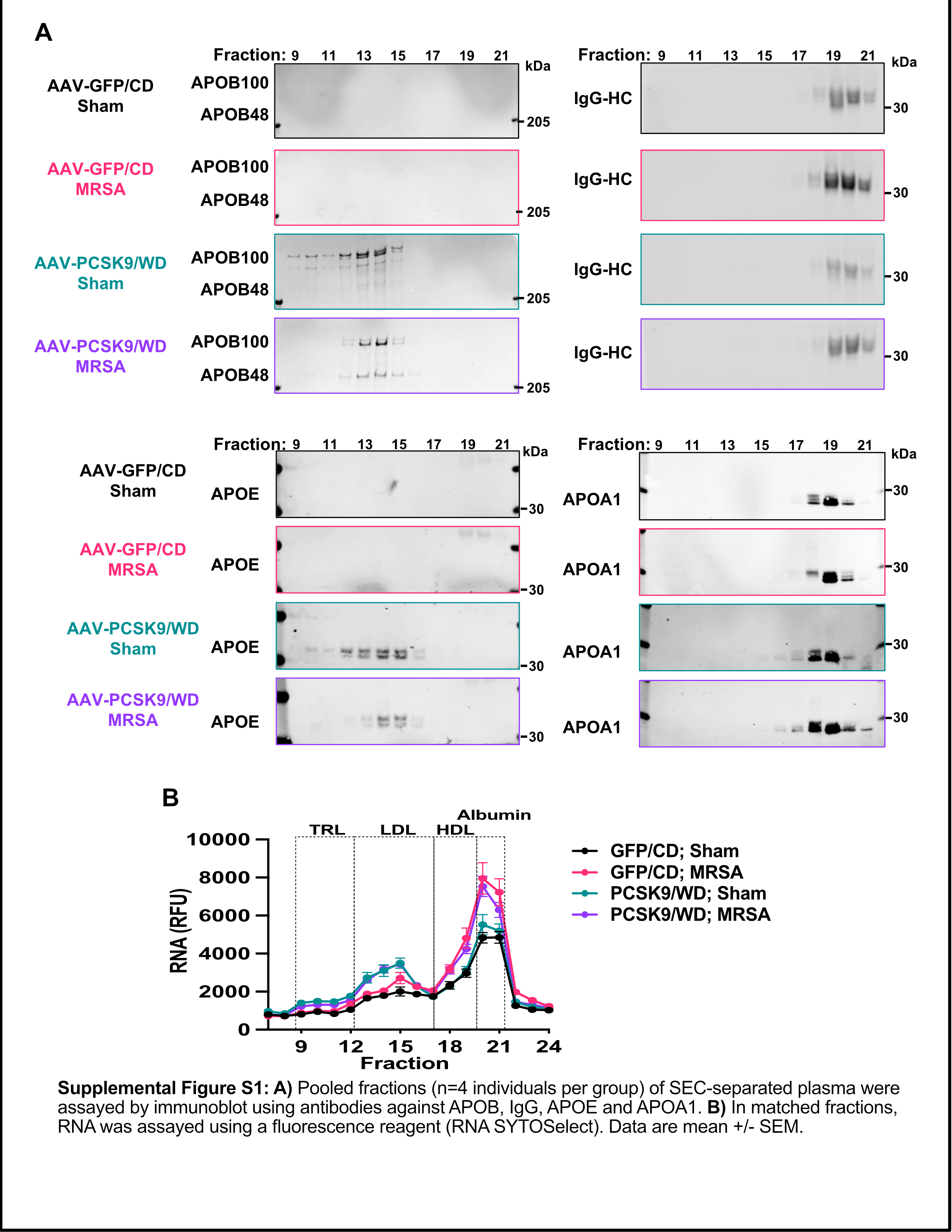

### Supplemental Figure S2

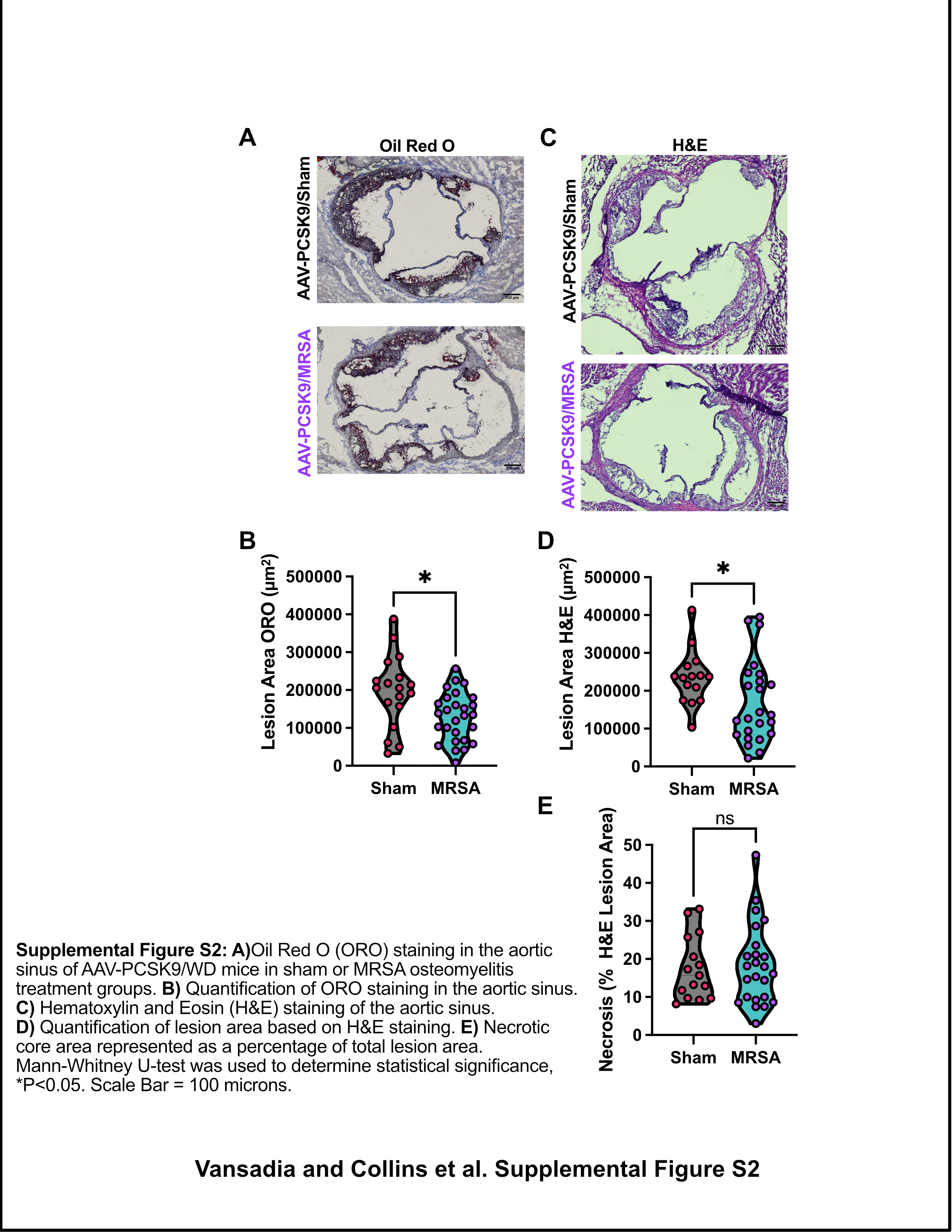

### Supplemental Figure S3

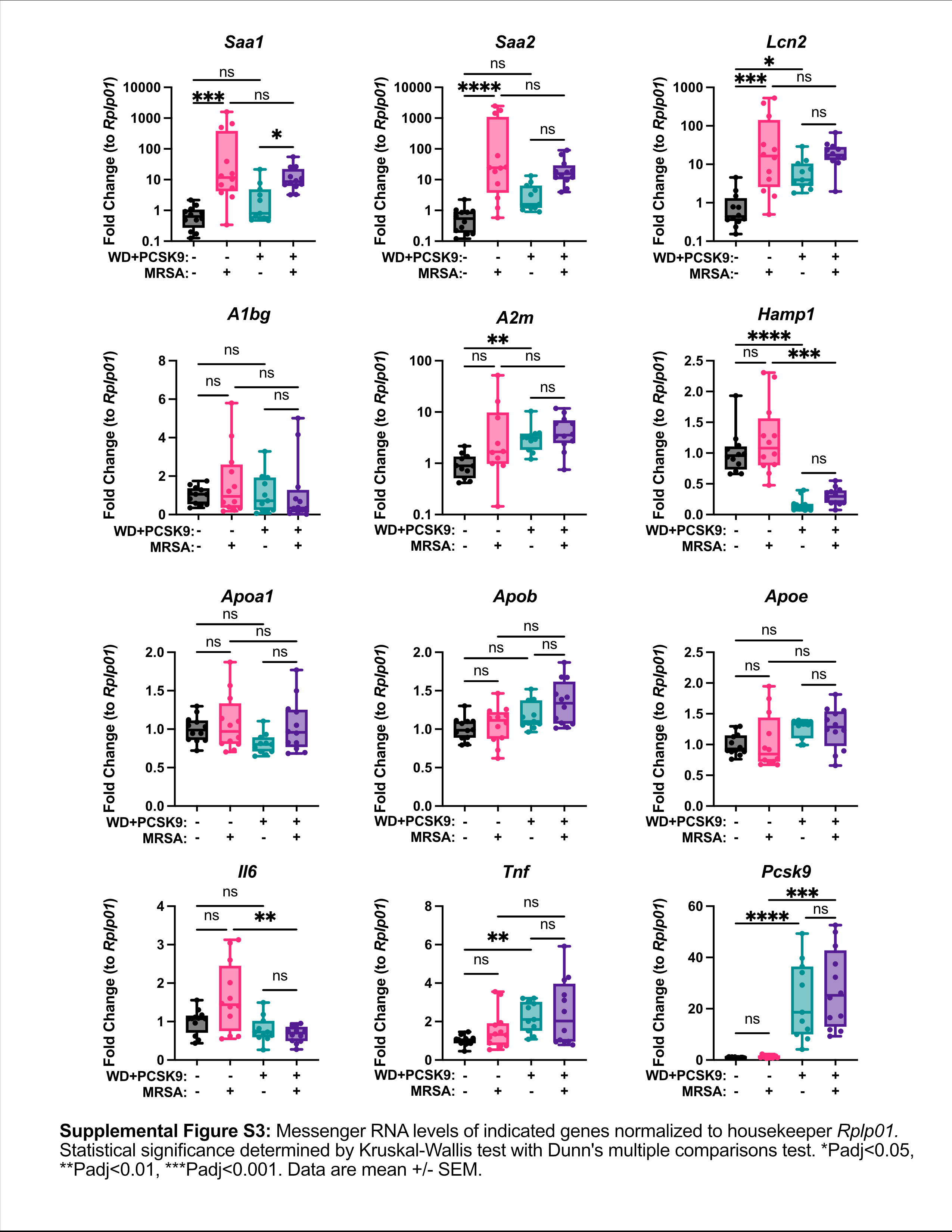
